# Using the Collaborative Cross to Dissect Immunological Mechanisms Governing Latent Infection, Granuloma Formation, and Neuroinvasive Disease Following Repeated Very Low-Dose Inhalation Exposure to Cryptococcus

**DOI:** 10.64898/2026.06.12.731915

**Authors:** Ivy M. Dambuza

## Abstract

**Background:** Decades of cryptococcal research have relied on a small number of hypervirulent laboratory strains, principally H99 and its derivatives, which cause rapid and lethal central nervous system infection and bypass the latent pulmonary phase that defines the natural history of human disease. As a result, the immune mechanisms that govern durable fungal containment in the lung, the reasons these mechanisms fail in genetically susceptible hosts, and whether the pulmonary immune response can engage the brain before any fungal cell reaches the central nervous system remain poorly understood. We address these gaps using a physiologically relevant repeated low-dose inhalation model in members of the Collaborative Cross and the BXD mouse panels.

**Methods:** A repeated low-dose intranasal exposure model, consisting of four doses of 50 *C. neoformans* UgCl223 cells over 18 days, was compared with a single primary challenge in C57BL/6J mice. Six Collaborative Cross founder strains (A/J, C57BL/6J, 129S1/SvImJ, NOD/ShiLtJ, NZO/H1LtJ, CAST/EiJ) and one BXD founder (DBA/2J) were assessed for fungal burden, granuloma architecture, and CD4 T cell polarisation. The neuroimmune axis was interrogated by exposing BV2 microglia to serum from infected mice with confirmed lung-restricted infection or disseminated disease.

**Results:** Repeated low-dose exposure, in contrast to single-dose primary challenge, was linked with enhanced fungal containment, elevated frequency of CD44^+^ CD4 T cells, expansion of regulatory T cells, and organised granuloma formation. Across seven genetically diverse strains, host genetic background determined disease outcome, spanning four orders of magnitude in lung fungal burden, from near-sterilising control in NZO mice to overt central nervous system dissemination with brain lesions in CAST mice. Granuloma architecture varied markedly across backgrounds in a pattern that mirrors human granuloma heterogeneity. The Th17 and regulatory T cell axis, rather than Th1 or Th2 polarisation, distinguished progressors from controllers. Critically, serum from mice with confirmed lung-restricted infection induced morphological microglial activation in a strain-dependent pattern associated with the disease spectrum, providing initial evidence supporting the existence of a lung-brain immune axis in cryptococcal disease.

**Conclusions:** These data demonstrate that the immune dynamics of latent cryptococcal infection are fundamentally distinct from those characterised in primary high-dose models. Host genetic background influences not only pulmonary containment of *Cryptococcus* but may also impact the magnitude of distal brain immune responses, potentially through mechanisms that extend beyond direct fungal dissemination to the central nervous system. These findings suggest that early pulmonary immune events may shape subsequent neuroimmune outcomes and identify a previously underappreciated lung-brain axis in cryptococcosis that warrants *in vivo* validation.

## Introduction

Cryptococcal meningitis affects approximately 194,000 people each year and causes an estimated 147,000 deaths, a case fatality of 75.8%, spanning both HIV/AIDS-associated and non-HIV immunodeficiency-related disease, predominantly in sub-Saharan Africa.^1^ Yet this burden represents only the catastrophic endpoint of a biological relationship that begins decades earlier in early childhood. Serological studies demonstrate that up to 70% of children in urban populations carry immunological evidence of prior *C. neoformans* exposure by age five,^2,3^ establishing that environmental inhalation of fungal spores is nearly universal. In immunocompetent individuals, the immune system contains the fungus within pulmonary granulomas without clinical symptoms, creating a state of latent infection that can persist for years before immune disruption triggers reactivation and dissemination.^3-6^ Despite substantial advances in defining antifungal immune pathways, a central unresolved question in cryptococcal pathogenesis is how most exposed hosts successfully contain *Cryptococcus* while others progress to pulmonary, disseminated, or central nervous system (CNS) disease.

Progress in answering this question has been constrained by a fundamental limitation in experimental models. Until recently, virtually all mechanistic understanding of *C. neoformans* immunity derived from studies using strain H99, first isolated on 14 February 1978 by Dr John Perfect at Duke University Medical Centre from a 28-year-old male with Hodgkin’s lymphoma and subsequently adopted as the universal laboratory reference strain.^7^ Decades of passage through laboratories and rabbit models generated divergent H99 lineages. The most widely distributed laboratory subcultures were later shown to belong to a hypervirulent lineage that harbours a 734-base-pair deletion in *SGF29*, a component of the SAGA histone acetyltransferase complex. This mutation substantially enhances virulence relative to the original clinical isolate.^8,9^ The congenic mating pair KN99α and KN99a, derived from this lineage, shares the same hypervirulent phenotype. When inoculated intranasally, these strains cause acute lethal meningoencephalitis within days to weeks and bypass the latent phase entirely. This has created a major blind spot, as the immune mechanisms that govern long-term fungal containment in the lung remained experimentally inaccessible using the strains that dominate the literature.^10^

Latency is not immunologically inactive but represents a dynamic equilibrium actively maintained by both host and pathogen. On the fungal side, *C. neoformans* deploys multiple strategies to persist within host tissues without triggering elimination. A subpopulation of dormant yeasts, characterised by low metabolic activity, increased autophagy, and regulated mitochondrial transcription, can enter a viable but non-culturable state that confers resistance to antifungal treatment and immune clearance.^11^ Reactivation of these cells is triggered by quorum-sensing molecules including vitamin B5, providing a mechanism for reactivation following immune disruption.^11^ In parallel, *C. neoformans* forms titan cells, which are dramatically enlarged yeast cells that can reach 100 μm in diameter during pulmonary infection.^12^ These cells are polyploid and resistant to phagocytosis because of their size, and they are proposed to contribute to latency by occupying granuloma niches in a state that limits immune recognition.^13^ Pathogen genetic variation further shapes the latency-versus-progression outcome. A recent genome-wide association study of ST93 clinical isolates identified four distinct in vivo disease phenotypes, including latent infection. Hypervirulence was associated with interferon gamma (IFN-γ)-dominated host responses and linked to single nucleotide polymorphisms (SNPs) in nine genes, including the inositol sensor *ITR4*, whose disruption recapitulated the hypervirulent phenotype.^14^ On the host side, CD4 T cells are the central mediators of cryptococcal containment. Their essential role is demonstrated by the strong association between CD4 depletion or functional impairment and the onset of progressive cryptococcal disease in conditions such as HIV and AIDS, glucocorticoid therapy, diabetes, or solid organ transplantation.^15,16^ However, most mechanistic characterisation of CD4-mediated immunity derives from primary infection models that use high-dose inocula of hypervirulent laboratory strains in single genetically uniform backgrounds. In this context, a dichotomy between protective Th1 responses and disease-permissive Th2 responses has been widely described. C57BL/6J mice, which mount a mixed Th1, Th2, and Th17 response to high-dose H99 infection, develop persistent pulmonary infection with an immunological stalemate, whereas strains with a stronger Th1 bias clear infection more effectively.^17,18^ IFN-γ is particularly protective in this setting, as it promotes macrophage fungicidal activity and supports granuloma integrity.^19^ Whether these dynamics operate during repeated low-dose exposure that mimics natural inhalation, where immune priming occurs gradually and regulatory circuits have time to develop, has not been investigated. Recent work on a mouse model of latent *C. neoformans* infection using the clinical isolate UgCl223 found that Tbet^+^ CD4 T cells predominate during latency, although they produce less IFN-γ.^20^ This finding diverges from the protective Th1 paradigm established in primary high-dose models and remains mechanistically unexplained. Paradoxically, IFN-γ-producing CD4 T cells are also implicated in cryptococcal neuropathology. In patients with cryptococcal post-infectious inflammatory response syndrome (PIIRS), a clinical syndrome defined by neurological deterioration despite microbiologically controlled infection, elevated cerebrospinal fluid (CSF) IFN-γ produced by antigen-specific CD4 T cells drives immune-mediated brain injury in the absence of active fungal replication.^21,22^ Mechanistically, brain-infiltrating fungal-specific CD4 T cells can drive inflammatory microglial proliferation through IFN-γ, generating a brain-resident immune cell population with poor fungicidal capacity that exacerbates CNS pathology even when fungal burden is controlled.^23^ These observations establish IFN-γ as a dual mediator in cryptococcal disease, supportive of protection during pulmonary infection yet capable of driving pathology within the brain. Critically, all existing data linking IFN-γ to CNS pathology derive from models of established brain infection. Whether IFN-γ or other inflammatory mediators generated during lung-restricted latent infection can engage the brain before any fungal cell arrives remains unknown. Host genetic variation profoundly shapes cryptococcal disease outcome, yet experimental genetics has been applied narrowly. Previous studies have mapped susceptibility loci on mouse chromosomes 1 and 9 using high-dose pulmonary infection^16^ and have shown that SJL mice recapitulate aspects of human resistance through differential immune activation.^24^ The Collaborative Cross (CC) is a reference population of recombinant inbred mouse lines derived from eight genetically diverse founder strains spanning four ancestral lineages, with genetic variation comparable to that seen in humans.^25,26^ This resource provides an unparalleled opportunity to dissect the host genetic architecture of infectious disease, yet it has not been applied to cryptococcal latency. Importantly, DBA/2J, one of the founder strains examined here, is also a founder of the BXD recombinant inbred panel, one of the most extensively characterised genetic reference populations in mouse biology. This panel contains fully annotated genotypes across thousands of markers as well as extensive phenotypic data available through GeneNetwork.^27,28^ Thus, both the CC and DBA/2J connect directly to existing quantitative trait mapping infrastructure and enable future high-resolution genetic analysis. Together, these findings establish that the transition from latent to progressive disease is shaped by both host and pathogen genetics. However, experimental models capable of dissecting this interaction have been lacking.

A critical gap in our understanding of cryptococcal disease is how pulmonary infection influences brain health. Clinical evidence shows that patients with asymptomatic cryptococcal antigenemia, defined by circulating fungal antigen in the absence of detectable CSF involvement, exhibit measurable neurocognitive deficits that are disproportionate to any detectable CNS fungal burden.^29^ This observation, reported in HIV-associated cohorts in Uganda, lacks a mechanistic explanation. It suggests that the immune response generated during lung-restricted infection may drive brain injury.

In this study, we establish a repeated low-dose intranasal exposure model using the clinical isolate UgCl223, which has been validated for latent infection in mice.^20,30^ This model recapitulates real-world environmental exposure through gradual immune priming followed by repeated challenge. Using this model across six CC and one BXD founder strains, we reveal a spectrum of disease outcomes determined by host genetics. We define the Th17 and regulatory T cell axis as the immunological tipping point between containment and progression. We also provide the first experimental evidence that circulating factors present in the serum of mice with lung-restricted *C. neoformans* infection can activate brain-resident microglia ex vivo, in the absence of central nervous system fungal dissemination. These findings reframe the immunopathogenesis of cryptococcal brain disease and identify a previously inaccessible window for early intervention.

## Materials and Methods

### Mouse Strains

Two infection models were established in female C57BL/6J mice aged 6–8 weeks, with 10–12 mice per group. Host genetic variation experiments used seven genetically diverse inbred founder strains, namely C57BL/6J, A/J, 129S, NOD, NZO, CAST (Collaborative Cross founders), and DBA/2J (BXD founder), with 3–5 mice per group. All strains were obtained from The Jackson Laboratory, USA.

### Infection Models

In the repeated low-dose exposure model, mice received three intranasal doses of 50 cells, consisting of ∼3 μm UgCl223 cells in 50 μl of phosphate-buffered saline (PBS), on days 0, 2, and 4. This was followed by a 14-day rest period to permit immune consolidation and granuloma formation, and then a single re-exposure on day 18 (Figure 1a). This schedule was designed to model sequential environmental encounters and to allow immunological priming and potential memory formation during the rest interval, before testing recall responses after re-exposure. Tissue harvest occurred at days 1, 7, and 32 post-exposure. The primary challenge model, used as a control, consisted of a single intranasal dose of 50 cells administered on day 0, with matched harvest time points (Figure 1b). UgCl223 is a Ugandan clinical isolate of the ST93 lineage that has been validated to cause persistent and controlled lung infection meeting criteria for latency in C57BL/6J mice.^20,30^ All experiments were conducted in accordance with the ethical review committee of the University of Exeter and UK Home Office regulations, under project licence number P6A6F95B5.

**Figure 1.**
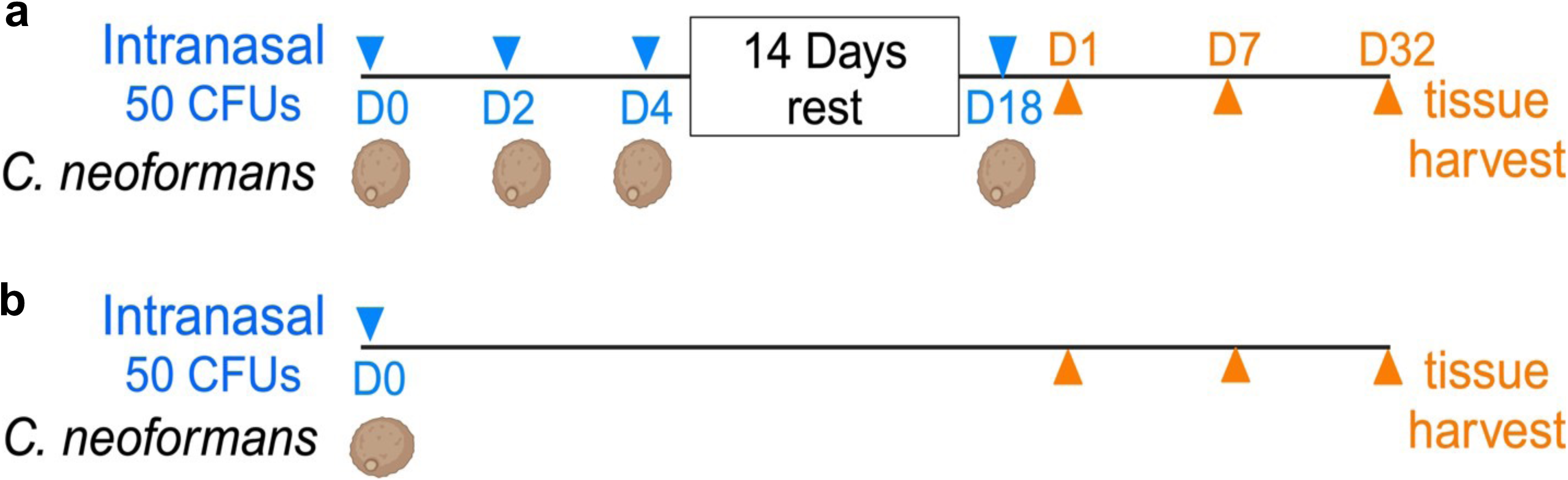
Very low dose repeat exposure model. To simulate constant environmental encounters with *C. neoformans*, **a)** female 6-8 weeks old C57BL/6J mice were challenged intranasally with 50 µl containing 50 cells of ∼3 µm UgCl223 cells at day 0, 2, 4, followed by 14 days rest then challenged again at day 18. **b)** Primary challenge model: single dose given at day 0. N=10-12 mice per group. Blue arrowheads = exposures, orange arrowheads = post-exposure

### Fungal Burden and Histology

At terminal harvest, lung, brain, liver, and spleen tissues were homogenised in 2 ml of PBS, serially diluted, and plated on yeast extract peptone dextrose agar. Colonies were enumerated after incubation at 30 degrees Celsius for 48 hours and reported as colony-forming units (CFU) per gram of tissue. For histological analysis, lungs were fixed in 10% formalin, embedded in paraffin, sectioned at 5 μm, and stained with haematoxylin and eosin (H&E). The day 32 timepoint was selected to capture established granuloma formation during latent infection, consistent with the phased granuloma progression described during latent cryptococcal infection.^31^

### Flow Cytometry

Lung-draining lymph nodes were processed into single-cell suspensions. CD4+ T cell subsets were identified by flow cytometry using antibodies against Tbet for Th1 cells, GATA3 for Th2 cells, RORγt for Th17 cells, Foxp3 for regulatory T cells, as well as activation (CD25) and memory (CD44) markers, at day 7 post-exposure. Data were acquired on a Cytek Aurora and analysed using FlowJo version 10.

### Ex Vivo Microglial Activation Assay

BV2 microglial cells were cultured in Dulbecco’s modified Eagle medium (DMEM) supplemented with 10% heat-inactivated serum collected from C57BL/6J, A/J, 129S, NOD, NZO, CAST, and DBA/2J mice at day 7 post-exposure, or from uninfected controls, for 24 hours. Cells were imaged at 20X magnification. Microglial morphological activation states were classified as resting or flat (red arrows), contracted lamellipodia or granulated (pink arrows), elongated or bipolar (yellow arrows), amoeboid (blue arrows), or ramified (green arrows), according to established criteria for morphological classification of microglial activation.^32,33^

### Statistical Analysis

Data were analysed using GraphPad Prism version 10. Two-group comparisons were performed using Student’s t-test. Multi-strain comparisons were conducted using two-way analysis of variance with Dunnett’s multiple comparison testing. A p value < 0.05 was considered statistically significant. Pooled data are presented as mean ± SEM.

## Results

### Repeated low-dose exposure establishes latency and drives an enhanced CD4 T cell response associated with fungal control

We first asked whether repeated exposure to *C. neoformans* produces immune outcomes that are qualitatively distinct from those observed following a single primary challenge. In the repeated exposure model, mice received three low-dose priming exposures followed by a 14-day rest interval designed to allow granuloma formation and immune consolidation, before re-challenge on day 18 (Figure 1a). At day 32, mice in the repeated exposure group exhibited a significant reduction in lung fungal burden compared with single-dose controls (p < 0.05; Figure 2a). Colony-forming units (CFUs) in the brain, liver, and spleen were below the limit of detection in the repeated exposure model, confirming lung-restricted infection and validating the model for studying latent, non-disseminating cryptococcal disease. Repeated exposure therefore generates a state that satisfies established criteria for latency, including persistent low-burden pulmonary infection, absence of systemic dissemination, organised granuloma formation, and CD4 T cell-linked containment.

**Figure 2.**
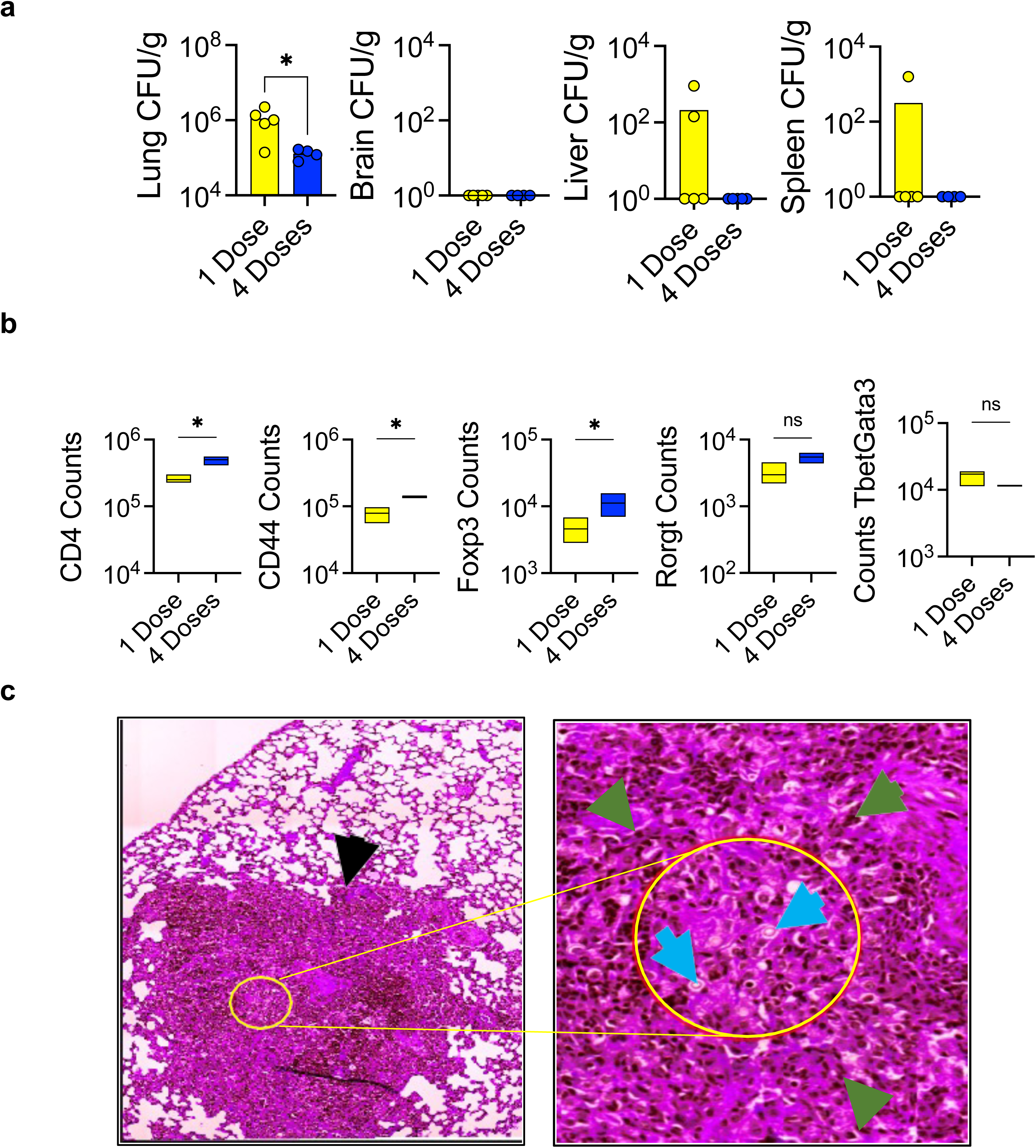
Improved capacity to control *C. neoformans* growth in the lungs of mice repeatedly exposed to very low doses of *C. neoformans* is associated with enhanced CD4 responses. **a)** Lung, brain, liver and spleen fungal burden determined at day 32 post exposure. **b)** Number of lung activated (CD44) CD4 cells and the phenotype of subsets assessed by transcription factor expression: Foxp3 (Tregs), RoRγt (Th17), Tbet (Th1), Gata3 (Th2). TbetGata3 refers to cells co-expressing both transcription factors. Single dose = blue bars/circles. Four doses = yellow bars/circles. **c)** Lung granuloma (black arrowhead) and *C. neoformans* in the centre (blue arrowheads) with granuloma immune cell types (green arrowheads) seen with H&E staining at day 32 post exposure. 40x magnification. N=4-5 mice per group, * p<0.05, *t* test

Improved fungal control was accompanied by selective expansion of activated pulmonary CD4 T cells (Figure 2b). Mice exposed repeatedly showed significantly increased total CD4 cell counts and higher frequencies of CD44^+^ activated CD4 T cells compared with single-dose controls (p < 0.05). Foxp3^+^ regulatory T cells were also significantly elevated (p < 0.05). Notably, RORγt^+^ and Tbet^+^GATA3^+^ populations did not differ significantly between groups. This profile, characterised by elevated activated CD4 T cells, is consistent with the establishment of antigen-experienced memory T cells together with immunoregulatory adaptation. These findings contrast with IFN-γ-dominated Th1 responses described in primary high-dose infection models using H99^17,18^ and instead align with observations that Tbet^+^ CD4 T cells in UgCl223-induced latency produce reduced IFN-γ.^20^ Together, these results suggest that protective immunity during true latency operates through a regulatory and memory axis distinct from the effector Th1 programme that drives clearance during acute primary infection.

The expansion of CD44^+^ and Foxp3^+^ CD4 populations, combined with improved fungal control after re-challenge, is consistent with the induction of immunological memory during the 14-day rest interval. Immunological memory is the adaptive immune system’s capacity to respond with accelerated vigour upon antigen re-exposure, and the survival of antigen-experienced CD4 T cells through contraction into a long-lived memory pool is a defining feature of this process.^34,35^

Histological examination confirmed the presence of discrete, well-organised granulomas containing *C. neoformans* at their centres (blue arrowheads) and surrounded by structured immune infiltrates (green arrowheads) in mice exposed repeatedly (Figure 2c). Formation of these granulomas correlated with improved fungal control and is consistent with the structured immune architecture required for long-term containment.

### Host genetic background determines disease trajectory

To test whether host genetic variation shapes the outcome of latent cryptococcal infection, Collaborative Cross founder strains (C57BL/6J, A/J, 129S, NOD, NZO, CAST) and the BXD founder DBA/2J were exposed to the repeated low-dose model and assessed over 28 days. Lung fungal burden at day 7 post-exposure spanned nearly four orders of magnitude across strains (Figure 3b). NZO mice, representing a controller phenotype, exhibited lung CFUs ∼2-log lower than C57BL/6J mice (p < 0.001), with no detectable brain infection (Figure 3c). In contrast, CAST mice were clear progressors, with lung burdens in the range of 10^7^–10^8^ CFUs per gram (p < 0.001 versus C57BL/6J) accompanied by substantial brain dissemination and grossly visible brain lesions. DBA/2J mice displayed a moderate progressor phenotype with brain dissemination, lung CFUs ranging from 10^3^–10^5^ colony-forming units per gram, defining a distinct intermediate phenotype. C57BL/6J, A/J, 129S, and NOD strains maintained a brain-negative status with intermediate lung burdens and therefore constitute a tolerant group. NZO mice showed a progressive increase in body weight to approximately 120 percent of baseline by day 28, reflecting a distinct metabolic and inflammatory profile associated with this controller strain. Because all animals received identical UgCl223 inocula, the observed spectrum of outcomes, which ranges from near-sterilising pulmonary control to lethal CNS dissemination, can be attributed entirely to host genetic variation. This is consistent with previously identified susceptibility loci on mouse chromosomes 1 and 9 mapped using high-dose infection models^16^ and with clinical observations in humans showing that sequence-type-matched isolates can produce markedly different outcomes depending on host genetic background.^14^

**Figure 3.**
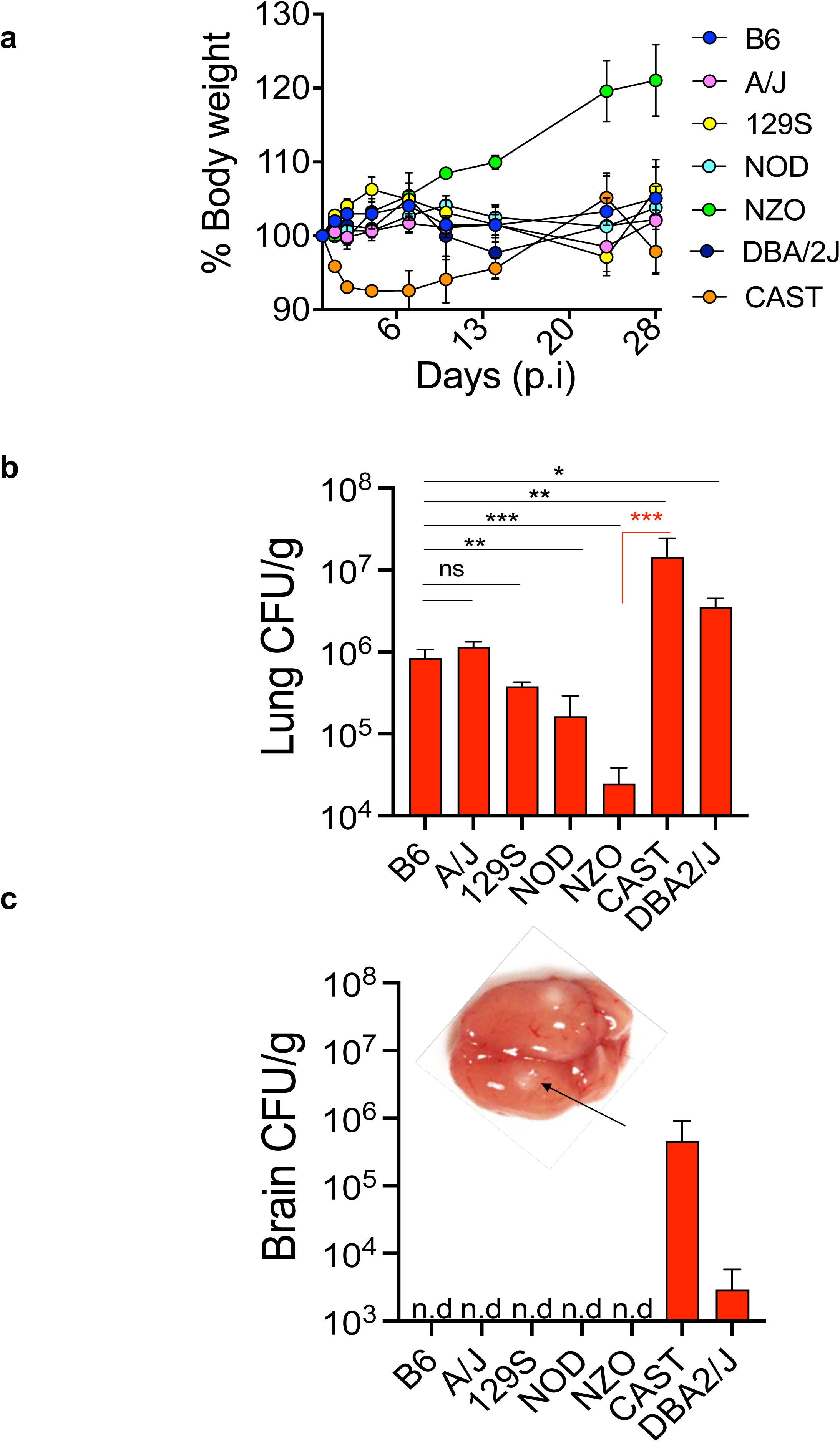
Host genetic variation impacts control of latent cryptococcosis, determining disease progression. Female 6-8 weeks B6, A/J, 129S, NOD, NZO, CAST and DBA/2J mice were exposed to repeat exposure model. **a)** Body weight changes. **b)** Lung and **c)** brain fungal burden determined day 7 post exposure. The Controller strain NZO has the least lung fungal burden (∼2-log difference compared to C57L/6 mice) and no brain infection, while the progressor strain, CAST, has the most dissemination. *Black arrow = brain lesions*. N=3-5 mice per group. * p<0.05, ** p<0.01, * p<0.001 Two-way ANOVA and Dunnett’s multiple comparison test.

The three-phenotype spectrum observed here, consisting of controller, tolerant, and progressor, closely corresponds to the *in vivo* virulence phenotypes of latent, non-central nervous system, central nervous system, and hypervirulent disease described using pathogen genetic variation.^14^ The convergence of host and pathogen genetic diversity on a shared disease spectrum provides complementary and independent support for the clinical heterogeneity of cryptococcal outcomes. It also establishes the Collaborative Cross and BXD founder strains as powerful and tractable systems for dissecting the immunogenetic determinants that define each disease state.

### Granuloma architecture is genetically determined and mirrors human clinical heterogeneity

Lung histology at day 7 post-exposure revealed striking strain-dependent variation in granuloma number, size, and architectural organisation (Figure 4). Similar to the controller strain NZO, NOD granulomas appeared irregular and flat. Among progressors, CAST lungs showed the most extensive pathology, with large, coalescing granulomatous masses, while DBA/2J lungs showed widely distributed prominent lesions. B6 and A/J showed moderate, well-circumscribed lesions; 129S showed larger, more numerous foci. Because all strains received identical inocula, inter-strain variability in granuloma morphology reflects host immune genetics rather than pathogen variation, directly mirroring the heterogeneity of granulomatous responses observed in human cryptococcal disease.^36^ These data suggest that effective latency requires not merely granuloma formation, but granuloma integrity, a distinction with direct therapeutic implications.

**Figure 4.**
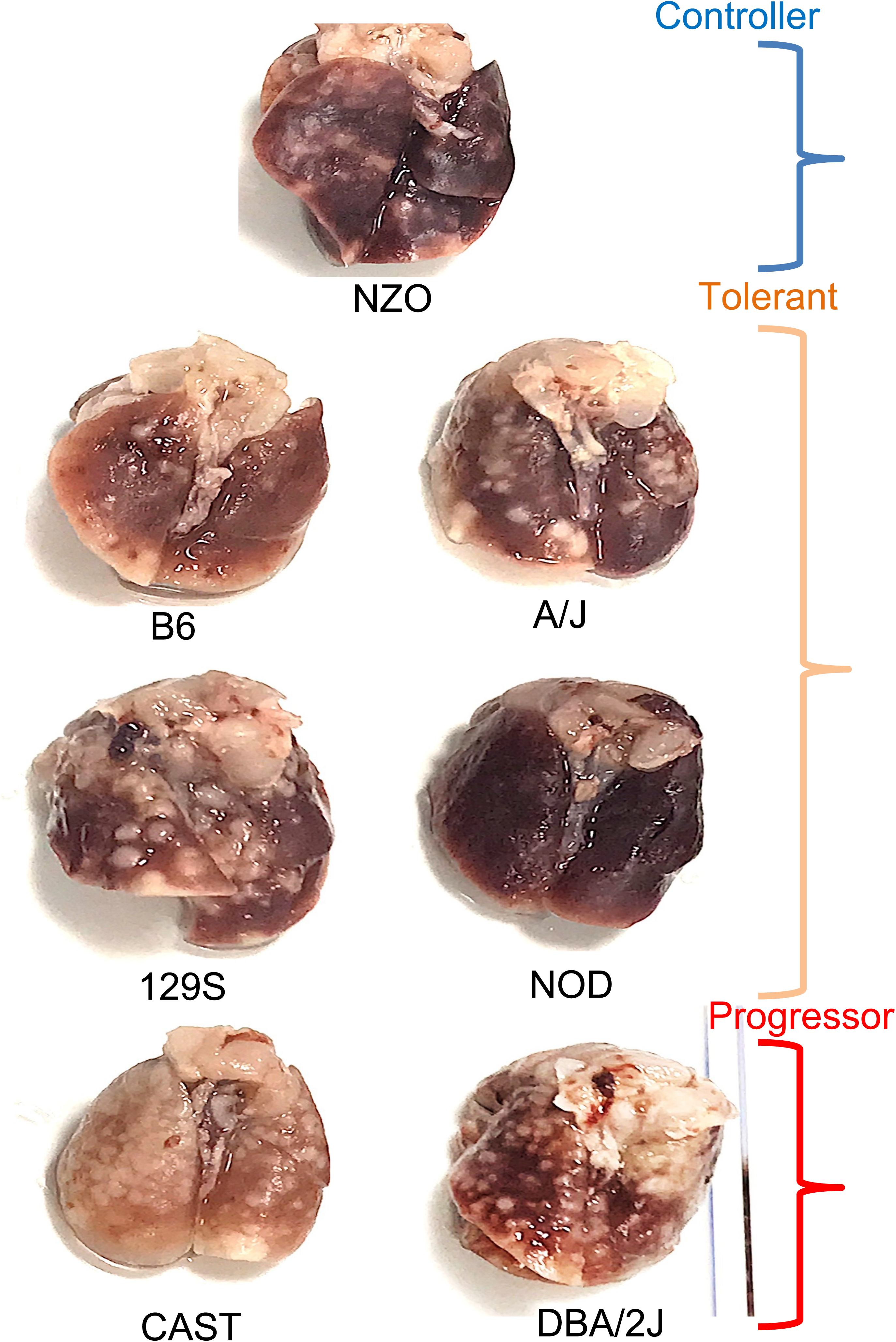
Host genetic variation regulates granuloma response. Lung representative images showing varied granuloma size lesions, number and shape, analyzed day 7 post exposure. N=3-5 mice per group. Variability in granuloma number and size mirrors the spectrum of granulomatous responses observed in humans. The variability observed using a single *C. neoformans* strain demonstrates that the host’s genetic make-up is a key driver of differences in disease outcome. Disease spectrum related to CFUs grouped together.

### The Th17-Treg axis distinguishes progression from containment

We next profiled CD4 T cell subsets in lung-draining lymph nodes across all seven strains to define the immunological correlates of the controller-progressor spectrum (Figure 5). Despite spanning four orders of magnitude in fungal burden, no significant inter-strain differences were observed in Th1 (Tbet^+^), Th2 (GATA3^+^), or activated (CD44^+^) CD4 populations (data not shown). This is a critical distinction from studies of primary high-dose *C. neoformans* infection with H99-derived strains, in which Th1/IFN-γ-dominated responses are the primary correlate of protection and IL-4/IL-13-driven Th2 polarisation drives permissive macrophage responses and fungal persistence.^17,18^ In the context of repeated low-dose exposure mimicking natural inhalation, Th1 and Th2 axes do not discriminate between controllers and progressors (data not shown), indicating that distinct immune mechanisms operate during latency. Instead, the Th17-Treg axis emerged as the genetically regulated discriminator of disease outcome. CAST progressors exhibited a significantly increased frequency of RORγt^+^ Th17 cells (p < 0.05) without a concomitant increase in Foxp3^+^ Tregs, compared to C57BL/6J. CD25 expression followed a pattern similar to RORγt. These findings implicate a dysregulated Th17 response as the immunological correlate of progression in the context of low-dose repeated exposure. While Th17 immunity is widely reported as protective in cryptococcal infection, promoting fungal clearance and macrophage activation in single-challenge, high-dose models,^37,38^ our finding that RORγt elevation marks the CAST progressor strain suggests a context-dependent role determined by exposure frequency and chronicity. A directly analogous phenomenon is well established in tuberculosis: IL-17 contributes to protective granuloma formation during acute infection, yet under repeated antigen exposure it becomes refractory to IFN-γ regulation and drives a hyperinflammatory, tissue-destructive response associated with disease progression.^39,40^ Our repeated low-dose model may therefore capture a similar transition, in which sustained Th17 activation reflects or contributes to failed containment rather than protective immunity.

**Figure 5.**
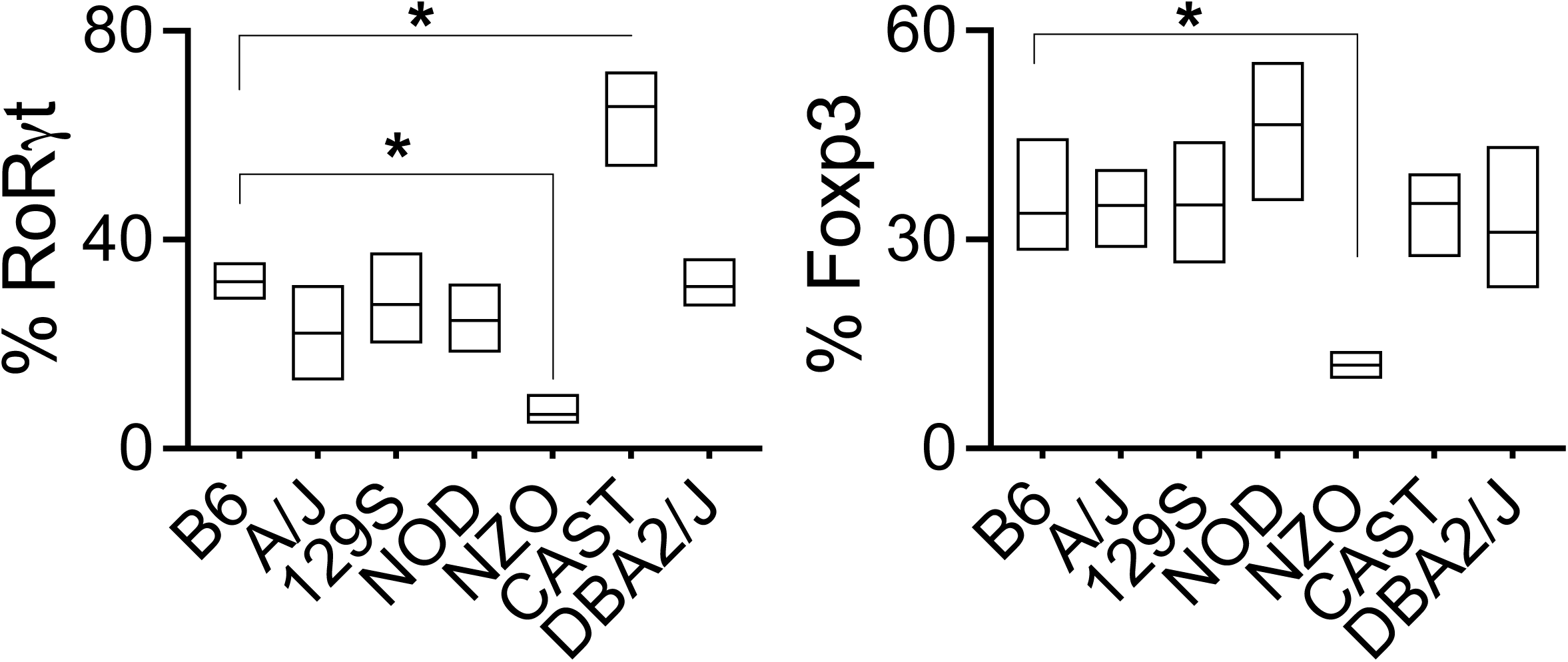
Host genetic background influences the Th17-Treg axis during latent cryptococcosis. Percentages of CD4+ T cells expressing T-bet (Th1), GATA3 (Th2), RORγt (Th17), Foxp3 (Tregs), CD25, and CD44 were quantified by flow cytometry in lung-draining lymph nodes at day 7 post exposure. CAST mice, which exhibited the greatest fungal dissemination, showed significantly elevated RORγt expression, suggesting that an exaggerated Th17 response may be associated with ineffective protective immunity. Relative to B6 mice, no differences were observed in CD44, T-bet, or GATA3 expression, whereas CD25 expression followed a pattern similar to RORγt (data not shown). Data represent 3–5 mice per group. Statistical analysis was performed using two-way ANOVA with Dunnett’s multiple-comparison test. *P < 0.05.

Principal component analysis of the transcription factors and markers measured (Tbet, GATA3, RORγt, Foxp3, CD25, and CD44) showed that NZO controllers separated from all other strains, but this separation was not explained by the transcription factors and markers measured here, suggesting that additional uncharacterised CD4 features are associated with their protective phenotype (PCA data not shown). Together, these data define the Th17-Treg balance as the critical immunological tipping point between durable latency and progressive disease in the context of physiologically relevant low-dose exposure.

### Circulating factors from lung-restricted infection activate brain-resident microglia: a lung-brain immune axis

The central conceptual question of this study, whether the pulmonary immune response to *C. neoformans* can engage the brain before any fungal cell arrives there, was addressed by exposing BV2 microglia to serum from each of the seven strains at day 7 post-exposure (Figure 6). Brain CFU was undetectable by plating in C57BL/6J, A/J, 129S, NOD, and NZO, confirming lung-restricted infection, and detected only in the progressors DBA/2J and CAST. Microglia cultured in serum from uninfected control mice maintained resting morphology (red arrows). In striking contrast, microglia cultured in serum from all seven infected strains exhibited activation-associated morphological changes in a strain-dependent pattern mirroring the disease spectrum. C57BL/6J serum induced contracted lamellipodia and granulated morphology (pink arrows); A/J and 129S induced elongated/bipolar morphology (yellow arrows); NOD and CAST serum induced pronounced amoeboid morphology (blue arrows: the signature of maximal pro-inflammatory activation); NZO and DBA/2J serum induced ramified morphology (green arrows: surveillant activation). Three findings are of particular mechanistic significance. First, NZO controller mice, with near-sterilising lung immunity and no detectable brain CFU, nonetheless produced serum that induced microglial morphological activation, indicating that the inflammatory mediators present have the potential to drive CNS immune engagement in the complete absence of fungal dissemination. Second, the strain-dependent gradient of microglial activation states, from moderate activation in controllers to maximal amoeboid transformation in progressors, mirrors the spectrum of pulmonary Th17-Treg polarisation and fungal burden, strongly implicating the systemic inflammatory milieu generated by the lung immune response as a potential driver. Third, CAST progressors, with the highest Th17 frequencies, greatest lung burden, and overt CNS dissemination, induced the most extreme microglial morphological response, suggesting that the same immune dysregulation that fails to contain the fungus in the lung also generates the most potent neuroimmune-activating circulating signal. These findings provide the first demonstration that serum from animals with lung-restricted fungal infection contains circulating factors capable of activating brain-resident immune cells *ex vivo*. The identity of the circulating factors responsible for microglial activation remains to be formally defined.

**Figure 6.**
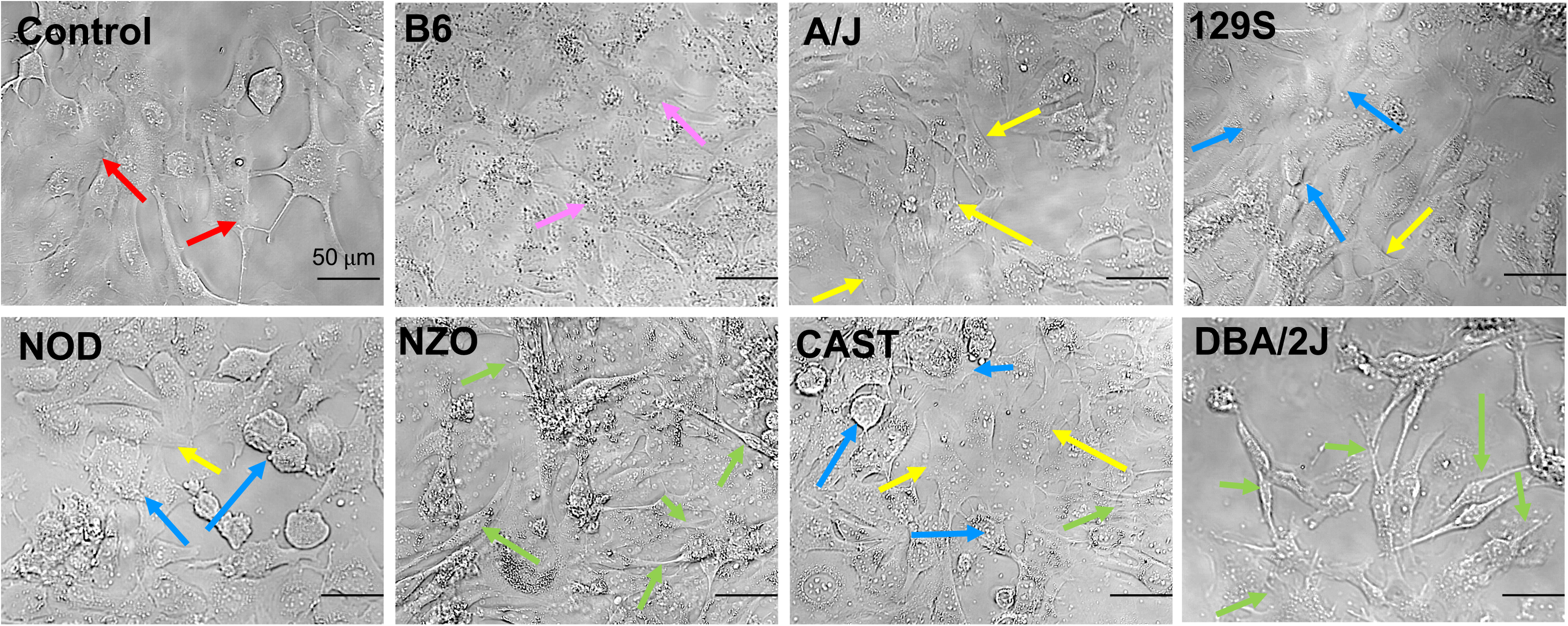
Factors present in the serum of mice repeatedly exposed to low-dose *C. neoformans* induce activation and stress-associated morphological changes in brain-resident cells. BV2 microglia cells were cultured in DMEM supplemented with serum from mice day 7 post-exposure or control (uninfected) for 24 hrs, then imaged on Olympus 20x magnification. N=4-5 mice per group. Arrows colour indicates microglia morphological changes. Red: resting/flat; Pink: contracted lamellipodia and granulated; Yellow: elongated/bipolar; Blue: amoeboid; Green: ramified morphology.

## Discussion

We report three principal findings. First, repeated low-dose *C. neoformans* exposure generates immune dynamics fundamentally distinct from those characterised in primary high-dose models: protective latency is maintained through a regulatory-memory CD4 axis featuring Treg expansion, not by the IFN-γ-dominated Th1 effector response that drives clearance in hypervirulent-strain infection. Second, host genetic background determines the entire spectrum of disease outcome, from near-sterilising containment to lethal CNS dissemination, across four orders of magnitude in fungal burden, with the Th17-Treg balance rather than Th1/Th2 polarisation distinguishing progressors from controllers in this context. Third, and most strikingly, serum from animals with confirmed lung-restricted infection activates brain-resident microglia *ex vivo* in a strain-dependent pattern mirroring the disease spectrum, providing the first experimental evidence consistent with a lung-brain immune axis in cryptococcal disease.

The experimental model established here addresses a critical and long-standing limitation of the field. Standard cryptococcal models using H99 and KN99, hypervirulent strains that cause acute lethal meningoencephalitis within days to weeks,^8,9^ make it impossible to study the immunology of latency, because the latent phase is bypassed entirely. Even clinical isolates at high inoculum doses produce outcomes more akin to acute infection than environmental exposure. The physiologically relevant scenario is one of repeated inhalation of very low spore numbers over time, during which the immune system is gradually primed, and memory circuits are established. Our model, using four doses of 50 UgCl223 cells over 18 days with a 14-day rest interval, approximates this process and generates a state of latency: lung-restricted infection, organised granulomas, and superior re-challenge control mediated by expanded CD44^+^ CD4 cells and Tregs. Others have reported repeated exposure at high dose (10^4^ cells), which generated an IL-4/IL-5 Th2/allergic response with worsening lung inflammation,^41^ highlighting the importance of physiologically relevant inoculum size in modelling natural exposures.

Our findings expand and contextualise the prior model of latent *C. neoformans* infection using a clinical isolate: Ding et al. used single-dose UgCl223 at 100 cells in C57BL/6J mice and identified Tbet^+^ CD4 cells as the predominant activated subset in lungs during latency, with decreased IFN-γ production compared to activated controls.^20^ Our repeated model generates an analogous state in C57BL/6J mice, with enhanced CD44^+^ activation without a Th17 or Th1/Th2 shift, alongside Treg expansion not captured in the single-dose system. Principal component analysis of the CD4 transcription factor landscape across all seven strains revealed that this picture is asymmetric: CAST progressors separate from all other strains along an axis driven by RORγt and CD25, co-directional markers indicating a coordinated, highly activated Th17 programme, providing a clear transcriptional signature of non-protective immunity. NZO controllers also separate as a distinct cluster, but not along this or any axis defined by the transcription factors measured (Tbet, GATA3, RORγt, and Foxp3), indicating that the features underlying their superior protection remain uncharacterised. The decreased IFN-γ in Ding’s model and our Treg expansion may reflect the same underlying phenomenon: active immune regulation during latency that dampens effector output to prevent immunopathology while preserving sufficient CD4 effector capacity for containment. This regulatory axis is fundamentally different from the protective Th1/IFN-γ programme documented in primary high-dose models.^17,18^ In progressor CAST mice, the emergence of a coordinated RORγt/CD25 Th17 signature, in the absence of a compensatory Treg increase, marks a shift away from this balanced regulatory state and correlates with failure to maintain containment. That protection in NZO mice cannot be explained by Tbet, GATA3, RORγt, or Foxp3 frequencies points to additional, as-yet-uncharacterised CD4 features, potentially functional, transcriptional, or metabolic, as determinants of durable latency, and identifies a priority target for future studies.

The CD44^+^ CD4 expansion and Foxp3^+^ Treg elevation after four doses are consistent with antigen-experienced memory T cells and memory Tregs.^34^ Enhanced fungal control after re-challenge compared to single-dose animals is consistent with a boosted recall response. Earlier work established that primary pulmonary *C. neoformans* infection can generate CD4-dependent protective memory: mice that had resolved a lung infection subsequently controlled the growth of blood-borne Cryptococcus yeast in the brain more rapidly than naive animals.^42^ Notably, however, that protection was Th1/IFN-γ-dependent and was demonstrated in a BALB/c-derived background following intravenous challenge, which is mechanistically distinct from the IFN-γ-independent regulatory-memory profile we observe during repeated low-dose latency. This distinction is significant given that Th1/IFN-γ responses, while protective in some contexts, drive microglial activation and CNS injury in others.^23^ Whether IFN-γ-independent memory is protective specifically because it avoids this pathological axis, and whether this is determined by host genetic background, infection dose, or chronicity, is a key question raised by our findings. The 14-day rest interval reflects the intermittent nature of real-world environmental exposure, where encounters with *C. neoformans* are separated in time, allowing immune responses to develop and consolidate between exposures rather than being driven by a single bolus challenge. Future studies in our laboratory will carry out adoptive transfer of CD44^+^ CD4 cells from repeatedly exposed mice into naive recipients followed by infection challenge, combined with TCR repertoire analysis to demonstrate clonal expansion of antigen-specific populations.

The extreme phenotypic contrast between NZO (controller) and CAST (progressor), spanning approximately four orders of magnitude in lung fungal burden and the presence or absence of CNS dissemination, in animals receiving an identical pathogen inoculum, provides direct evidence that host genetic background is a primary, tractable determinant of cryptococcal disease outcome. This magnitude of phenotypic separation between CC founder strains is precisely what makes the Collaborative Cross such a powerful resource for genetic dissection: the recombinant inbred CC lines, each carrying a unique mosaic of alleles from these eight founders including NZO and CAST, segregate the genetic determinants underlying this contrast across a panel with genetic diversity comparable to the human population.^25,26^ Future work extending our repeated low-dose model to the CC lines will enable quantitative trait locus (QTL) mapping of (i) lung fungal burden and containment efficiency, (ii) Th17-Treg balance, and (iii) serum-mediated neuroimmune activation, testing directly whether the same loci that govern fungal containment also govern the magnitude of the lung-brain immune signal, or whether these traits are genetically separable. DBA/2J offers an additional and complementary route to genetic resolution: it is a founder of the BXD recombinant inbred panel,^27,28^ the most extensively characterised genetic reference population in mouse biology, comprising over 100 lines derived from B6 x DBA/2J crosses with full genotype annotation and integrated phenotypic databases available through GeneNetwork (www.genenetwork.org). The progressor phenotype of DBA/2J, with higher lung burden than C57BL/6J strain, establishes a tractable phenotypic contrast for QTL mapping within the existing BXD infrastructure. Collectively, the CC and BXD panels provide complementary genetic platforms to generate the first comprehensive map of host determinants governing cryptococcal latency and the lung-brain immune axis. The resulting insights would have direct translational relevance for the identification of human genetic risk variants.

Indeed, host genetic mapping of cryptococcal susceptibility has precedent, though using high-dose models. Carroll et al. performed genome-wide QTL analysis in 435 (C3H/HeN × CBA/J)F2 hybrids using lung fungal burden as the phenotype, identifying two significant loci on chromosomes 1 and 9.^16^ Notably, the resistant CBA/J parental strain in that study was characterised by greater neutrophilia and higher pulmonary IFN-γ, CXCL10, and IL-17 compared to the susceptible C3H/HeN strain, an association between IL-17/Th17 signalling and resistance that appears, on the surface, to contrast with our finding that RORγt elevation marks the progressor CAST strain. We note, however, that Carroll et al. used a high-dose primary infection model, in which IL-17 may contribute to acute neutrophil-mediated clearance, whereas our repeated low-dose model probes a chronic latency setting in which sustained RORγt/CD25 co-expression may instead reflect a dysregulated, non-resolving Th17 state. This dose-and chronicity-dependent reversal of Th17 function has clear precedent in tuberculosis, where IL-17 is protective during acute infection but pathological under repeated antigen exposure.^39,40^ Independently, pathogen genetic variation also shapes the latency-versus-progression outcome: Jackson et al. identified four *in vivo* disease phenotypes including latent infection across a panel of closely related ST93 clinical isolates, with hypervirulence associated with an IFN-γ-dominated host response and mapped to SNPs in nine genes including the inositol sensor *ITR4*.^14^ That host genetic background (this study, and Carroll et al.^16^) and pathogen genetic background^14^ each independently produce a comparable spectrum of outcomes, from latent containment to disseminated disease, suggests that disease trajectory reflects the combined genetic contributions of both host and pathogen, a perspective that has been largely inaccessible using standard hypervirulent laboratory strains in genetically uniform hosts. However, neither prior QTL study addressed latency specifically, nor the CD4 T cell programmes that actively maintain it. Our finding that NZO controllers cannot be distinguished from other strains by Tbet, GATA3, RORγt, or Foxp3 frequencies underscores this gap: the CD4-mediated mechanisms that maintain durable latency may be genetically and mechanistically distinct from those governing resistance to acute high-burden infection, and remain entirely unmapped.

The *ex vivo* microglial activation data raise mechanistic questions about the nature of the circulating factors responsible. In PIIRS, CSF IFN-γ produced by antigen-specific CD4 T cells drives immune-mediated brain injury despite negative fungal cultures,^21,22^ demonstrating that IFN-γ alone, without active fungal replication, is sufficient to cause neurological harm. Others have shown that brain-infiltrating IFN-γ-producing CD4 T cells drive inflammatory microglia proliferation, generating a brain immune cell population with poor fungicidal capacity that exacerbates CNS pathology.^23^ These data establish the IFN-γ-microglia axis as one of the mediators of CNS injury. The identity of the circulating factor(s) responsible for the microglial activation we observe when using serum from our low-dose repeated exposure model, characterised by an enhanced memory CD4 response, remains undefined. In C57BL/6J, A/J, 129S, NOD, and NZO mice, brain CFU was undetectable (whole-brain homogenate plating), yet serum from these animals was sufficient to activate microglia *ex vivo*, indicating that this activation can occur independently of fungal presence and reflects the immune response to the lung-restricted infection. DBA/2J and CAST mice are the exceptions: with confirmed brain dissemination, their serum reflects an immune response that includes active CNS infection. Nonetheless, CAST, which showed the highest RORγt^+^ Th17 frequencies in the lung, raises the possibility that the same dysregulated Th17 programme may be a candidate contributor to the circulating factors. Formal identification requires multiplex profiling of infection serum across all strains, cytokine neutralisation in the *ex vivo* assay, and *in vivo* confirmation of microglial activation state during lung-restricted infection, a programme underway in our laboratory.

Study limitations are recognised. The microglial activation data derive from a BV2 *in vitro* model using bulk serum, which does not reproduce the complexity of the *in vivo* blood-brain barrier environment. Whether morphological changes translate to functional shifts in microglial inflammatory state requires in *vivo* confirmation. The identity of the circulating neuroimmune-activating factors, whether cytokines, fungal polysaccharide antigen, extracellular vesicles, or other circulating mediators, is undefined. Finally, the sex of the animals used (female) was chosen for consistency; sex-specific immune differences in cryptococcal infection have not been examined in this context and represent an important future variable.

## Conclusions

Using a novel repeated low-dose exposure model that recapitulates natural environmental encounters with *C. neoformans*, we demonstrate that the immune dynamics of cryptococcal latency are fundamentally distinct from those characterised in primary high-dose infection models employing hypervirulent laboratory strains. This model provides a tractable experimental platform for investigating the immunological mechanisms governing fungal containment, latency, and disease progression under conditions that more closely reflect natural exposure. Protective latency is linked with a CD4 T-cell programme that remains to be defined. Host genetic background shapes the full spectrum of disease outcomes across four orders of magnitude in fungal burden, with variation in the Th17-Treg balance emerging as a stronger correlate of outcome than classical Th1/Th2 polarisation. Most significantly, we provide experimental evidence that serum from animals with lung-restricted cryptococcal infection can activate brain-resident microglia *ex vivo* in the absence of detectable CNS fungal dissemination, with both the magnitude and character of microglial activation determined by host genetic background. Collectively, these findings support a model in which pulmonary immune responses during latent cryptococcal infection may influence distal neuroimmune activation. They further suggest that mechanisms contributing to cryptococcal neuropathogenesis could be initiated during the pulmonary phase of infection, well before overt CNS disease becomes clinically apparent. If validated in vivo, this concept would redefine the therapeutic window in cryptococcal disease, shifting attention from established meningitis to earlier pulmonary infection, when immune responses may remain amenable to intervention and the development of irreversible neurological injury could potentially be prevented.

## Author Contributions

I.M.D. conceived the study, designed and performed all experiments, analysed data, and wrote the manuscript.

## Acknowledgements

We thank the staff of the animal facilities at the University of Exeter for the care and support of our animals. We acknowledge funding from the MRC Centre for Medical Mycology at the University of Exeter (MR/N006364/2 and MR/V033417/1) and the NIHR Exeter Biomedical Research Centre (NIHR203320). Additional work may have been undertaken by the University of Exeter Biological Services Unit. The views expressed are those of the author and not necessarily those of the NIHR or the Department of Health and Social Care. For the purpose of open access, the author has applied a CC BY public copyright licence to any Author Accepted Manuscript version arising from this submission.

## Competing Interests

The author declares no competing interests.

## Notes

### Competing Interest Statement

The authors have declared no competing interest.

